# Accurate assembly of the olive baboon (*Papio anubis*) genome using long-­read and Hi-C data

**DOI:** 10.1101/678771

**Authors:** Sanjit Singh Batra, Michal Levy-Sakin, Jacqueline Robinson, Joseph Guillory, Steffen Durinck, Pui-Yan Kwok, Laura A. Cox, Somasekar Seshagiri, Yun S. Song, Jeffrey D. Wall

## Abstract

Besides macaques, baboons are the most commonly used nonhuman primate in biomedical research. Despite this importance, the genomic resources for baboons are quite limited. In particular, the current baboon reference genome Panu_3.0 is a highly fragmented, reference-guided (i.e., not fully *de novo*) assembly, and its poor quality inhibits our ability to conduct downstream genomic analyses. Here we present a truly *de novo* genome assembly of the olive baboon (*Papio anubis*) that uses data from several recently developed single-molecule technologies. Our assembly, Panubis1.0, has an N50 contig size of ~1.46 Mb (as opposed to 139 Kb for Panu_3.0), and has single scaffolds that span each of the 20 autosomes and the X chromosome. We highlight multiple lines of evidence (including Bionano Genomics data, pedigree linkage information, and linkage disequilibrium data) suggesting that there are several large assembly errors in Panu_3.0, which have been corrected in Panubis1.0.

## INTRODUCTION

Baboons are ground-living monkeys native to Africa and the Arabian Peninsula. Due to their relatively large size, abundance and omnivorous diet, baboons have increasingly become a major biomedical model system (reviewed in [1]). Baboon research has been facilitated by the creation (in 1960) and maintenance of a large, pedigreed, well-phenotyped baboon colony at the Southwest National Primate Research Center (SNPRC) and an ability to control the environment of subjects in ways that are obviously not possible in human biomedical studies. For example, baboons have been used to study the effect of diet on cholesterol and triglyceride levels in controlled experiments where all food consumption is completely controlled [2] [3] [4]. In recent years, linkage studies in baboons have helped identify genetic regions affecting a wide range of phenotypes, such as cholesterol levels [5] [6], estrogen levels [7], craniofacial measurements [8], bone density [9] [10] and lipoprotein metabolism [11]. In addition, studies have also documented that the genetic architecture of complex traits in baboons can be directly informative about analogous traits in humans (e.g.,[10] [12]).

The success of these and other studies have been mediated in part by recent advances in molecular genetics technologies. In particular, the ability to cheaply genotype and/or sequence samples of interest has led to a revolution in genetic studies of the associations between genotype and phenotype. While human genetic studies now routinely include the analyses of whole-genome sequence data from many thousands of samples (e.g., [13] [14] [15] [16] [17]), comparable studies in model organisms have lagged far behind. Part of the reason for this is the lack of genetic resources in non-human species. Large, international projects such as the Human Genome Project [18] [19], International HapMap Project [20] [21] [22] and the 1000 Genomes Project [23] [24] [25] have provided baseline information on sequences and genetic variation, and subsequent human genetic studies have utilized this background information.

The first published baboon genome assembly was from a yellow baboon [26]. This assembly used a combination of Illumina paired-end and Illumina mate-pair sequence data (with mean library insert sizes ranging from 175 bp to 14 Kbp) to produce a highly fragmented assembly with contig N50 of 29 Kbp and scaffold N50 of 887 Kbp. The public olive baboon assembly, Panu_3.0, suffers from the same problem of having small contigs and scaffolds (contig N50 of 139 Kbp and *de novo* scaffold N50 of 586 Kbp) [27]. The authors of the public olive baboon assembly chose to distribute a reference-guided assembly with scaffolds mapped onto rhesus (*Macaca mulatta*) chromosomes. As a consequence, any syntenic differences between rhesus and baboon will result in large-scale assembly errors in Panu_3.0. One additional drawback of this baboon genome assembly was its informal embargo from 2008 to 2019 under the guidelines of the Fort Lauderdale agreement. Hence, its influence on scientific research has been negligible.

In this project, we focus on providing a high-quality, *de novo* genome assembly for olive baboon (*Papio anubis*), which we call Panubis1.0, with the hope that this resource will enable future high-resolution genotype-phenotype studies. Unlike previous baboon genome assembly efforts, we use a combination of three recently developed technologies (from 10x Genomics linked-reads, Oxford Nanopore long reads, and Hi-C) to increase the long-range contiguity of our assembly. These newly developed technologies enable us to generate assemblies where the autosomes (and the X chromosome) are each spanned by a single scaffold at a cost that is orders of magnitude cheaper than the Panu_3.0 assembly. We also verify that most of the large-scale syntenic differences between our Panubis1.0 and Panu_3.0 are due to errors in the public assembly rather than our own. Our assembly is available for scientific use without any restrictions.

## RESULTS

### Genome Assembly

The main strength of our approach is in combining data from multiple platforms (10x Genomics linked-reads, Oxford Nanopore long-reads, Illumina paired-end short-reads, and Hi-C), which have complementary advantages. Figure 1 describes our assembly strategy. We began by assembling 10x Genomics reads generated with their Chromium system (average depth ~60x) using the SUPERNOVA assembler (version 1.1) [28], which yielded an assembly with a contig N50 of ~84 kb and a scaffold N50 of ~15.7 Mb (Table 1). The gap lengths between the contigs in a scaffold obtained by assembling 10x linked-reads are arbitrary [29]. Hence, in order to leverage the Oxford Nanopore long-reads for gap-closing, we split the 10X scaffolds at every stretch of non-zero N’s to obtain a collection of contigs.

**Table 1.**
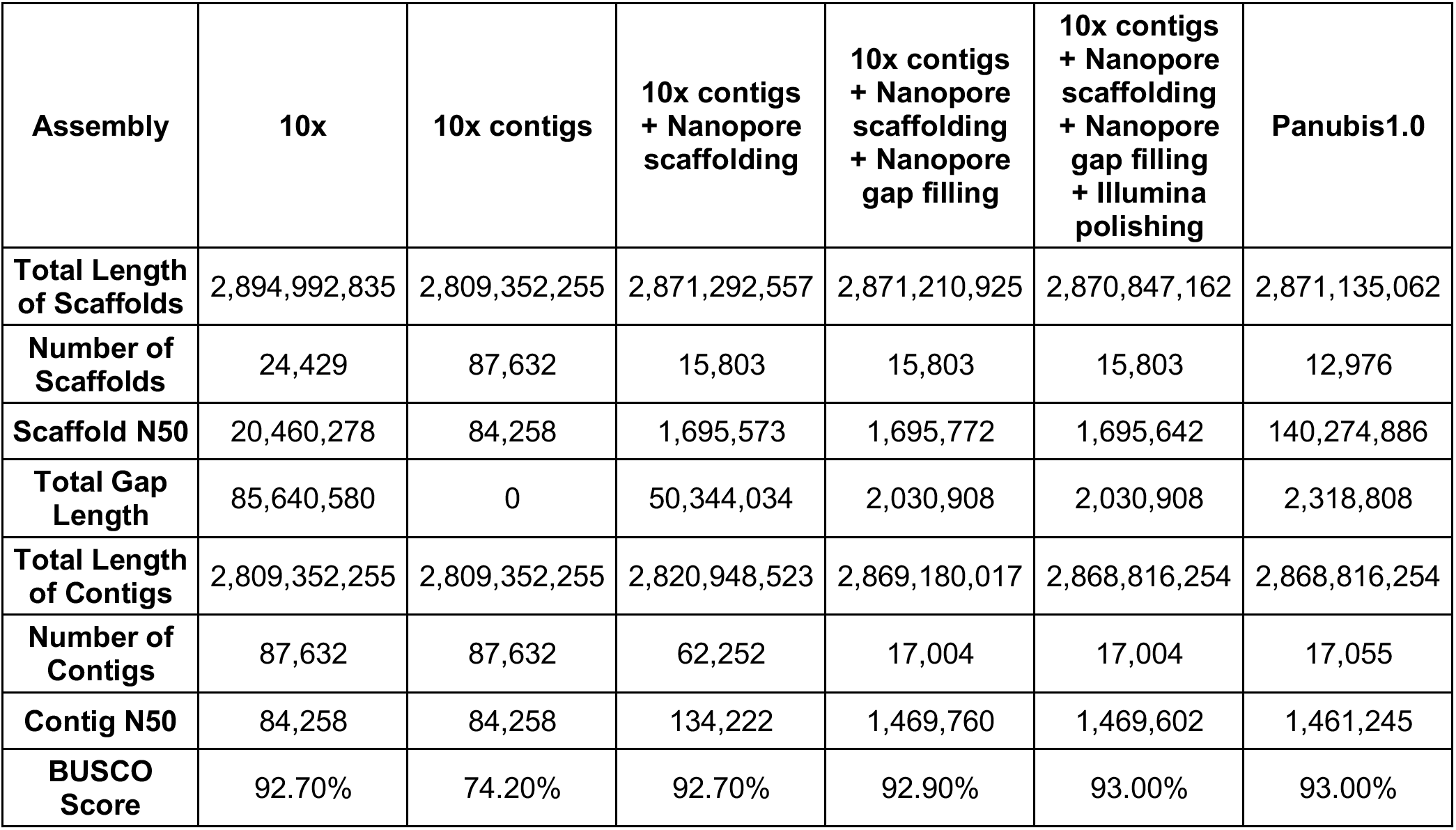
Assembly statistics for each step of the adopted assembly strategy. Total Length of Scaffolds is the sum of lengths of scaffolds (including A, C, G, T and N) in each scaffold. Total Gap Length is the total number of N’s in the assembly. Total Length of Contigs is the sum of the number of sequenced base pairs (including only A, C, G and T) in each scaffold. BUSCO provides a way of measuring the presence of genes conserved in mammals [37]. Since BUSCO reports complete genes and fragmented genes, the BUSCO Score is the fraction of complete mammalian BUSCO genes found in the assembly.

**Figure 1.**
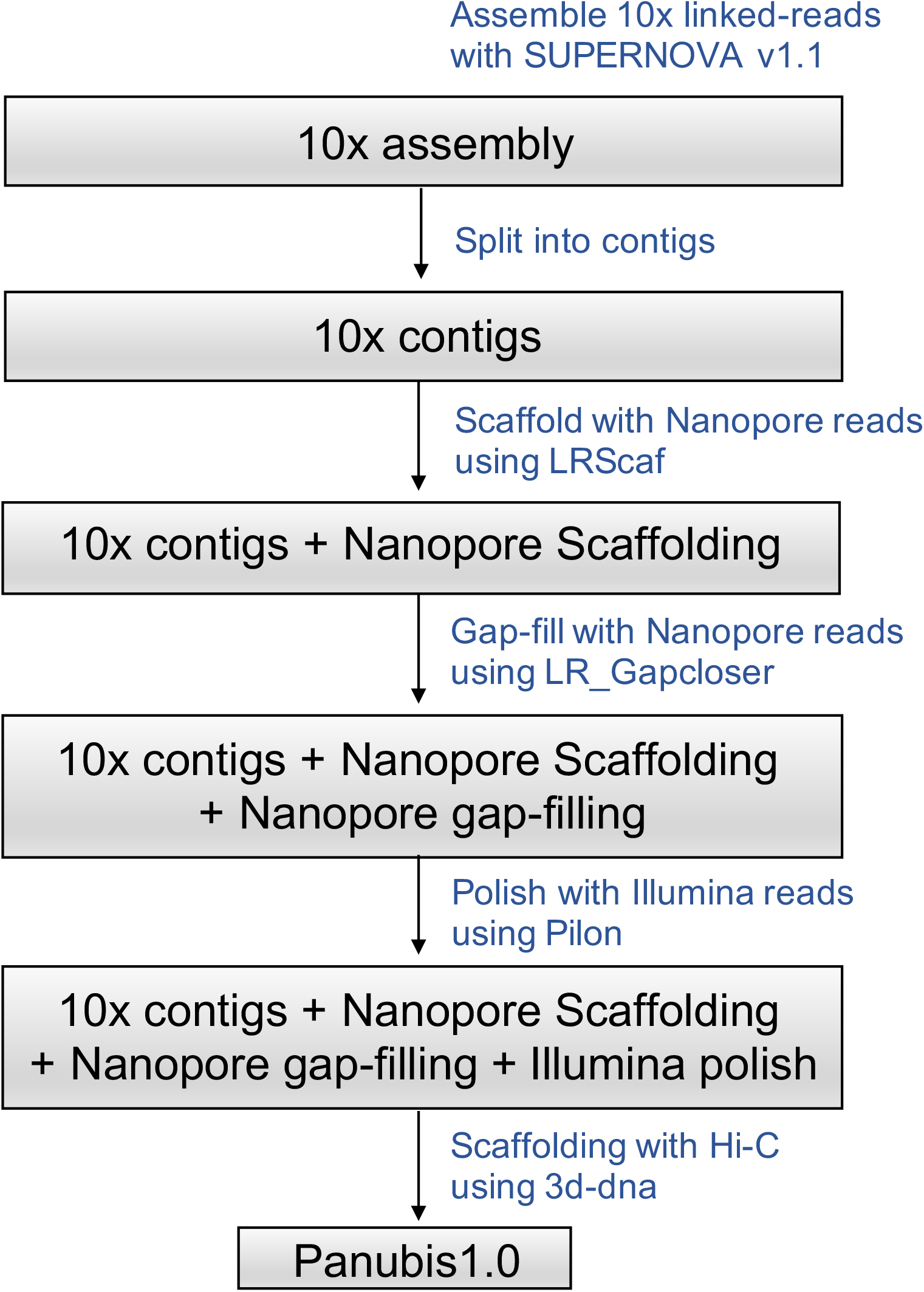
Illustration of our genome assembly strategy.

We scaffolded the resulting contigs with Oxford Nanopore long-reads (average depth ~15X) using the LR_Scaf [30] scaffolding method. This resulted in an assembly with a contig N50 of ~134 kb and a scaffold N50 of ~1.69 Mb (Table 1). These resulting scaffolds are more amenable to gap-closing, because the gap lengths (number of Ns between two consecutive contigs) are estimated by long-reads that span each gap and align to the flanking regions of that gap.

Upon performing gap-closing with the same set of Oxford Nanopore long-reads using LR_Gapcloser [31], we obtained an assembly with a contig N50 of ~1.47 Mb and a scaffold N50 of ~1.69 Mb (Table 1). Note that this increase in contig N50 of ~84Kb from the 10x Genomics linked-read assembly, to a contig N50 of ~1.47 Mb, would not have been possible if we had simply performed gap-closing with the Oxford Nanopore long reads directly on the 10x-based assembly without first splitting it into its constituent contigs. Finally, we polished the resulting assembly by aligning Illumina paired-end reads (average depth ~60X in PE150 reads) using Pilon [32].

In order to scaffold the resulting assembly with Hi-C data, we first set aside scaffolds shorter than 50 kb, which comprised only ~1.8% of the total sequence base pairs. This was done because Hi-C based scaffolding is more reliable for longer scaffolds, since there are more Hi-C reads aligning to longer scaffolds. We then ordered and oriented the remaining scaffolds using the 3D *de novo* assembly (3d-dna) pipeline [33] using ~15X Hi-C data generated by Phase Genomics [34]. Finally, we manually corrected misassemblies in the resulting Hi-C based assembly by visualizing the Hi-C reads aligned to the assembly, using Juicebox Assembly Tools [35], following the strategy described in [36]. Figure 2 shows Hi-C reads aligned to the resulting assembly with the blue squares on the diagonal representing chromosomes.

**Figure 2.**
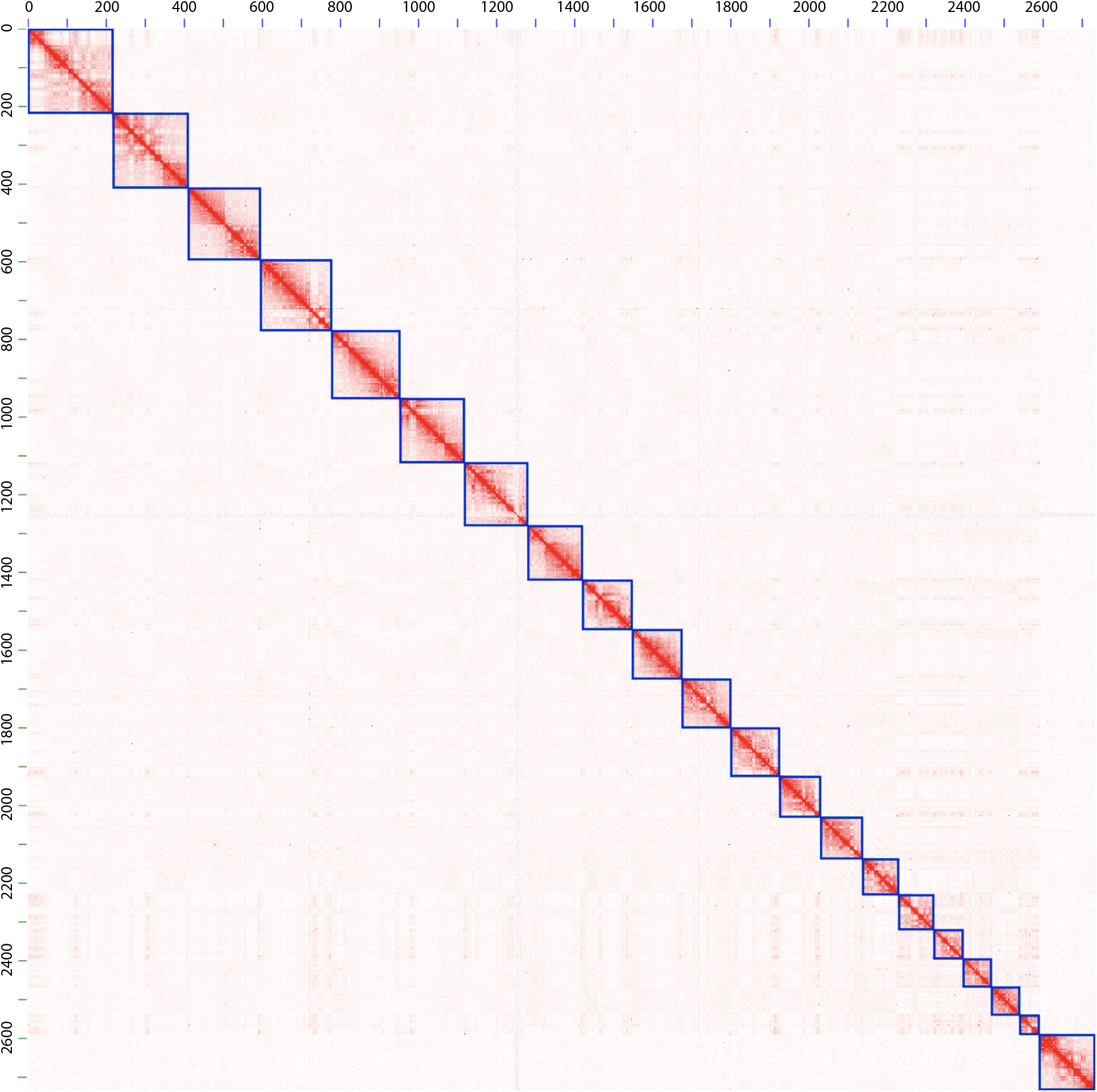
The Hi-C map of our Panubis1.0 genome. Each blue square on the diagonal represents a chromosome-length scaffold. Autosomes are listed first, ordered by size, and the last square corresponds to the X chromosome. The axes are labelled in units of megabases.

The resulting *Papio anubis* genome assembly, which we name Panubis1.0, contains ~2.87 Gb of sequenced base pairs (non-N base pairs) and 2.3 Mb (<0.1%) of gaps (N’s). Single scaffolds spanning the 20 autosomes and the X chromosome together contain 95.14% (~2.73 Gb) of the sequenced base pairs. We number the autosomes as chr1 to chr20, in decreasing order of the scaffold length, so some chromosome numbers in our convention are different from Panu_3.0’s numbering. We note that Panubis1.0 has a contig N50 of 1.46 Mb, which is a greater than ten-fold improvement over the contig N50 (~139 kb) of the Panu_3.0 assembly. As a result, Panubis1.0 contains five times fewer scaffolds (12,976 scaffolds with a scaffold N50 of ~140 Mb) compared to the Panu_3.0 assembly (63,235 scaffolds with a scaffold N50 of ~586 Kb); see Table 2 for a further comparison of the two assemblies. Gene completion analysis of the assembly using BUSCO v2 and the odb9 Mammalia ortholog dataset [37] suggests that Panubis1.0 contains 93.00% complete genes, comparable to the Panu_3.0 assembly.

**Table 2.** Comparison of Panubis1.0 with Panu_3.0 assemblies.

| **Assembly** | **Panubis1.0** | **Panu_3.0** |
| --- | --- | --- |
| **Total Length of Scaffolds** | 2,871,135,062 | 2,959,373,024 |
| **Number of Scaffolds** | 12,976 | 63,235 |
| **Scaffold N50** | 140,274,886 | 585,721 |
| **Total Gap Length** | 2,318,808 | 22,434,732 |
| **Total Length of Contigs** | 2,868,816,254 | 2,937,001,527 |
| **Number of Contigs** | 17,055 | 122,216 |
| **Contig N50** | 1,461,245 | 138,819 |
| **BUSCO Score** | 93.00% | 93.40% |

### Y chromosome assembly

The Hi-C scaffolding with 3d-dna yielded an ~8 Mb scaffold that putatively represents part of the baboon Y chromosome. Since, rhesus macaque is the phylogenetically closest species to baboons which has a chromosome-scale assembly, we aligned this putative baboon Y chromosome scaffold with the rhesus macaque Y chromosome (Figure 3). We observed a substantial amount of synteny between the putative baboon Y and the rhesus Y, comparable to what is observed between the chimpanzee Y and the human Y chromosomes. (For comparison, genetic divergence between baboon and rhesus is similar to human – chimpanzee divergence [38].) The observed breaks in synteny are consistent with the well-documented high rate of chromosomal rearrangements on mammalian Y chromosomes [39].

**Figure 3.**
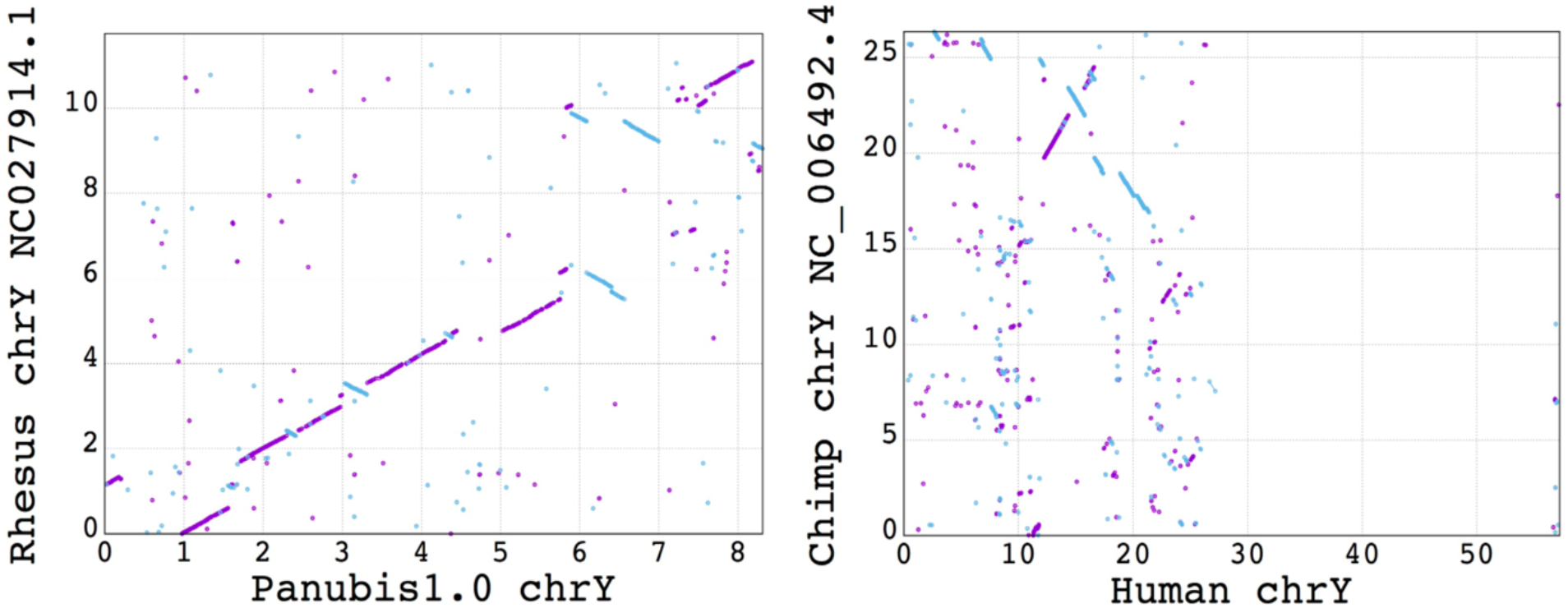
Dotplots showing chromosome Y synteny. A dotplot between rhesus chromosome Y and Panubis1.0 putative chromosome Y is shown on the left, while a dotplot between the chimpanzee chromosome Y and the human chromosome Y is shown on the right. The axes labels are in units of megabases. The phylogenetic distance between baboon and rhesus macaque is similar to that between human and chimpanzee. Hence, the broadly conserved synteny between the rhesus and baboon putative chromosome Y as compared to the synteny between the chimp and human chromosome Y, suggests that the scaffold representing the putative chromosome Y in the Panubis1.0 assembly is indeed capturing at least a large part of chromosome Y.

### Comparisons with the publicly available Panu_3.0 assembly

Figure 4 presents a dotplot between the chromosomes of the Panubis1.0 and the Panu_3.0 assemblies. There are chromosomes with large differences between the two assemblies and these differences are evident even in the chromosome-scale dotplots. Table 3 presents a list of large (>100 Kb) differences between the Panubis1.0 and Panu_3.0 assemblies. We used several orthogonal sources of information to assess whether these were errors in our Panubis1.0 assembly or in the Panu_3.0 assembly. These included Bionano Genomics optical maps obtained from the same individual used for generating Panubis1.0, linkage information from a pedigree of baboons that were all sequenced to high coverage, and linkage-disequilibrium information from 24 unrelated olive baboons from the SNPRC pedigreed baboon colony. We manually examined each break in synteny between Panubis1.0 and Panu_3.0 to determine whether these independent sources of evidence supported one assembly over the other (summarized in Table 3). Overall, in 11 out of 12 large syntenic differences between Panubis1.0 and Panu_3.0, at least one of these independent sources provided evidence that the Panubis1.0 assembly is correct. These independent sources of evidence make it overwhelmingly likely that the Panubis1.0 assembly provides the correct order and orientation for the sequence. For the remaining large syntenic difference, it is difficult to conclude which one of Panubis1.0 and Panu_3.0 is correct. An example of the nature of this evidence is displayed in Figure 5, which shows that the region starting at ~29.38 Mb and ending at ~44.71 Mb on scaffold NC_018167.2 in Panu_3.0 is inverted relative to the Panubis1.0 assembly. We provide additional information in support of the Panubis1.0 assembly from several other regions in Supplementary Figures S1-S5.

**Table 3.** Potential large (>100 Kb) assembly errors in Panu_3.0, ordered by size. Note that a ‘no’ in the ‘Linkage support’ or ‘LDhelmet support’ columns is inconclusive, and should not be interpreted as support for the Panu_3.0 assembly being correct. ^1^Unable to determine whether linkage and LDhelmet provide support at the end breakpoint due to a lack of synteny between Panu_2.0 and Panu_3.0 ^2^Panu_2.0 assembly appears to be correct ^3^BNG maps do not support a translocation with these breakpoints. However, they do support a potential large SV at the starting breakpoint ^4^BNG maps support the presence of a large SV, which may be a translocation ^5^Linkage data suggests a potential polymorphic inversion (in 16413) partially overlapping with this interval

| Panu_3.0 chromosome | Panu_3.0 Start | Panu_3.0 End | Panu_2.0 Start | Panu_2.0 End | Type | Linkage support | BNG support | LDhelmet support |
| --- | --- | --- | --- | --- | --- | --- | --- | --- |
| NC_018164.2 | 88.05 | 104.99 | 87.61 | 104.98 | Inv | start<sup>1</sup> | yes | unknown<sup>1</sup> |
| NC_018167.2 | 29.38 | 44.71 | 29.25 | 44.53 | Inv | start + end | yes | start + end |
| NC_018156.2 | 4.04 | 8.67 | 4.18 | 8.63 | Inv | no | yes<sup>2</sup> | no |
| NC_018162.2 | 82.42 | 86.47 | 81.91 | 84.01 | Trans | start + end | no<sup>3</sup> | no |
| NC_018166.2 | 104.28 | 108.05 | 103.66 | 107.44 | Inv | no | yes | no |
| NC_018165.2 | 15.93 | 19.48 | 15.85 | 19.40 | Inv | no | no | no |
| NC_018166.2 | 96.94 | 100.12 | 96.39 | 99.54 | Trans | start + end | yes<sup>4</sup> | start + end |
| NC_018160.2 | 36.05 | 36.75 | 35.88 | 36.55 | Trans | no | yes<sup>4</sup> | start |
| NC_018163.2 | 23.19 | 23.66 | 0 | 0.47 | Trans | no | yes<sup>2</sup> | no |
| NC_018164.2 | 4.05 | 4.49 | 3.99 | 4.45 | Trans | no<sup>5</sup> | yes | no |
| NC_018165.2 | 100.91 | 101.18 | 100.31 | 100.59 | Trans | no | yes | no |
| NC_018152.2 | 166.73 | 166.89 | 169.86 | 170.10 | Trans | start + end | yes | end |

**Figure 4.**
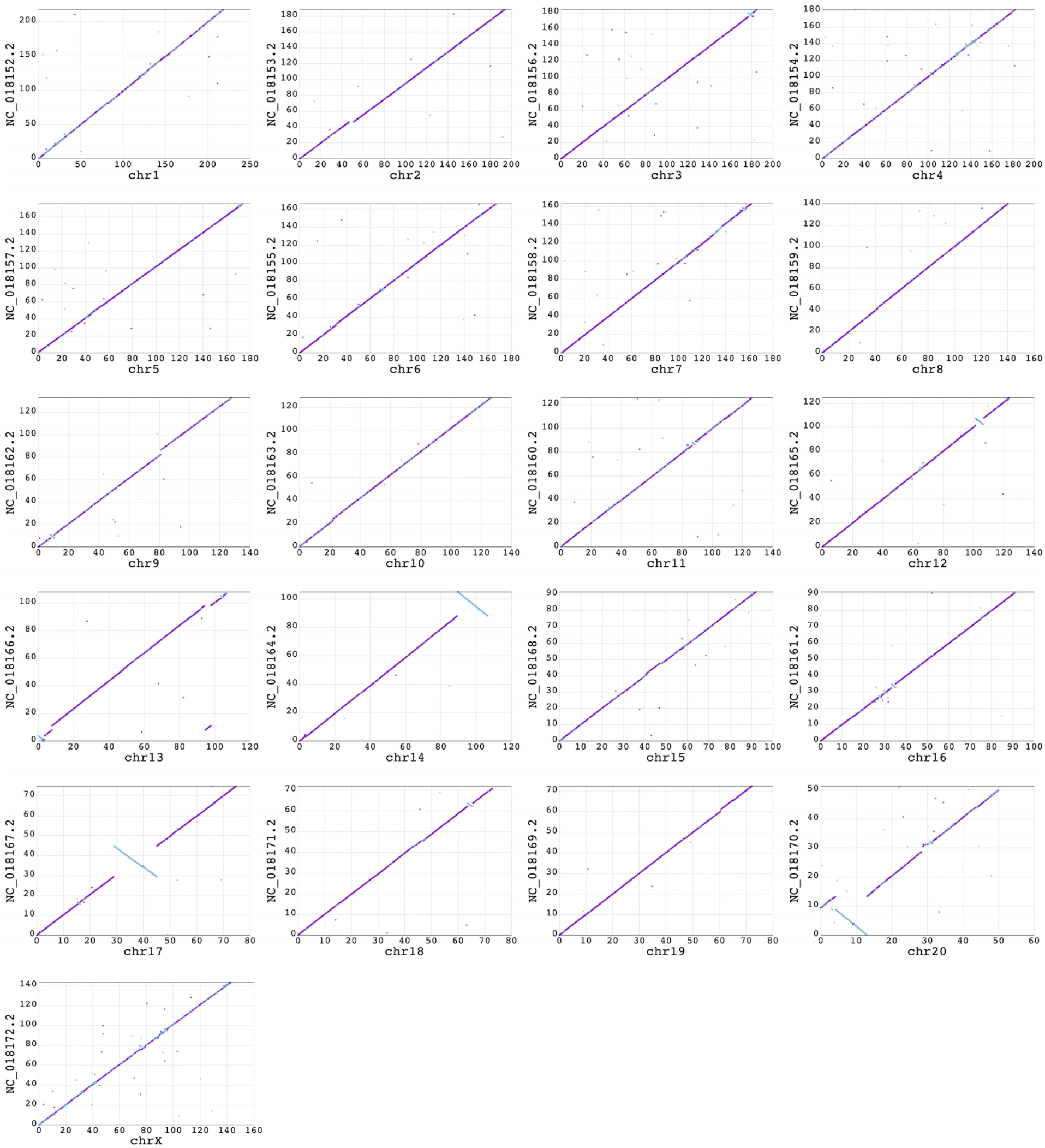
Dotplots showing alignment of Panu_3.0 reference-assisted chromosomes vs. Panubis1.0 chromosome-length scaffolds. The Panu_3.0 assembly is shown on the Y-axis and the Panubis1.0 assembly is shown on the X-axis. Each dot represents the position of a syntenic block between the two assemblies as determined by the nucmer alignment. The color of the dot reflects the orientation of the individual alignments (purple indicates consistent orientation and blue indicates inconsistent orientation). The dotplots illustrate that there are chromosomes containing large inversions and translocations in the Panu_3.0 assembly with respect to the Panubis1.0 assembly.

**Figure 5.**
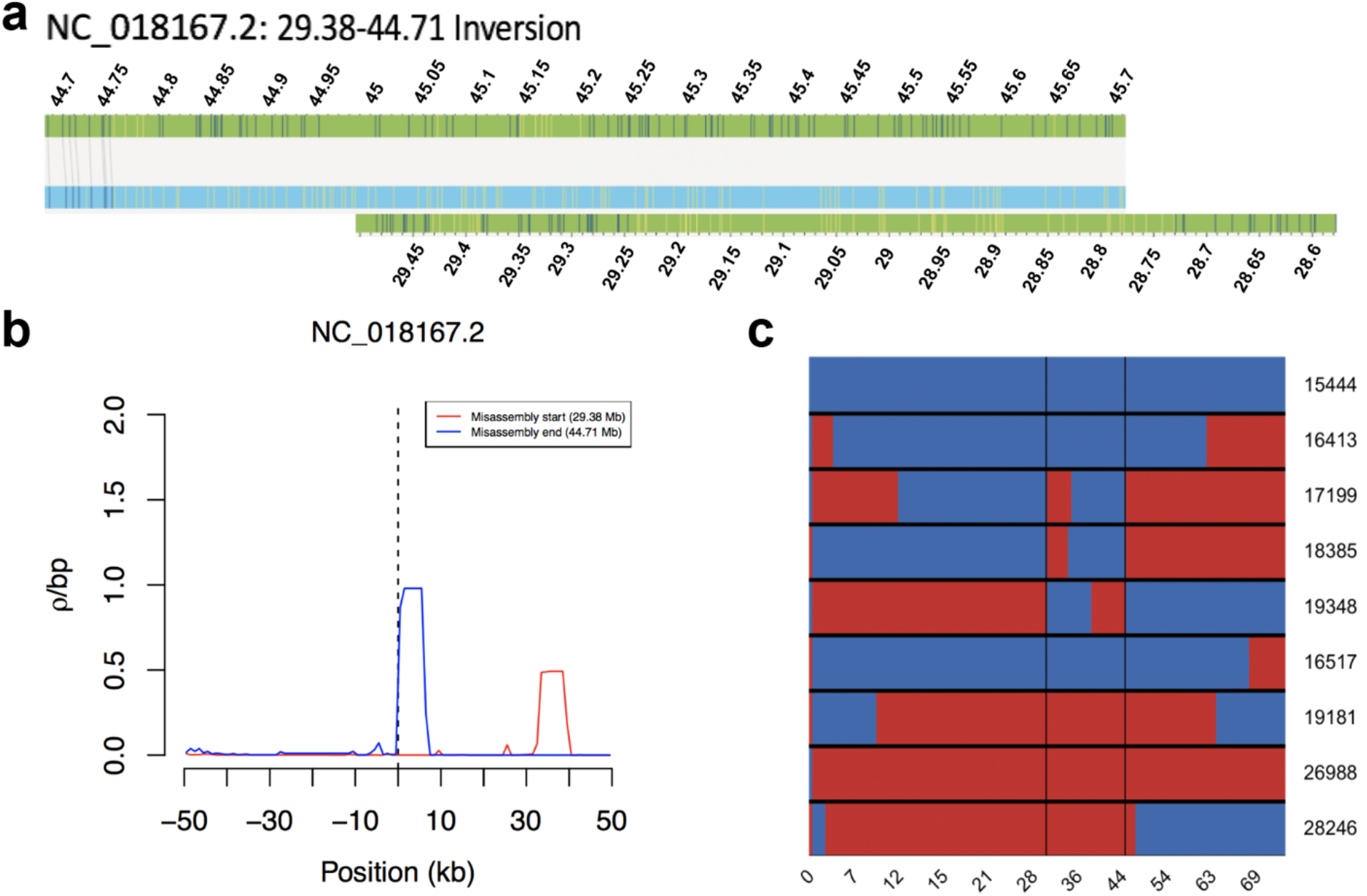
Evidence for misassembly on chromosome NC_018167.2 in Panu_3.0. **a)** Bionano optical map alignment to the Panu_3.0 assembly demonstrates there is an inversion on chromosome NC_018167.2 beginning at ~29.38 Mb and ending at ~44.71 Mb. **b)** Estimates of the population recombination rate ρ near the potential synteny breaks of the inversion identified on chromosome NC_018167.2. **c)** Shown on the x-axis is positions along chromosome NC_018167.2 in Panu_3.0 where each row represents one of the 9 offsprings of sire 10173. Switches between red and blue within a row represent a recombination event. The two vertical black lines represent locations where three or more recombinations occur at the same locus indicating a potential misassembly.

## DISCUSSION

The development and commercialization of new technologies by companies such as Illumina, 10x Genomics, Bionano Genomics, Dovetail Genomics and Phase Genomics has enabled researchers to cheaply generate fully *de novo* genome assemblies with high scaffold contiguity (e.g., [40]; [41]; [33]; [36]; [42]). When used in combination with long-read sequences (e.g., from Oxford Nanopore or Pacific Biosciences), these technologies can produce high-quality genome assemblies at a fraction of the cost of traditional clone library based approaches (e.g., [41]; [43]). In this context, our assembly Panubis1.0 provides a 10-fold increase in contig N50 size and a 240-fold increase in scaffold N50 size relative to Panu_3.0 at less than 1% of the reagent cost. The contiguity of this assembly will be especially useful for future studies where knowing the genomic location is important (e.g., hybridization or recombination studies).

One natural question that arises with any new genome assembly is how one assesses that an assembly is *‘correct’*. Indeed, some of the recently published Hi-C based assemblies have not provided any corroborating evidence supporting their assemblies (e.g., [44]). Here, we used three independent sources to provide evidence that 11 out of 12 large syntenic differences are correct in our new baboon assembly (Panubis1.0) relative to the previous assembly Panu_3.0 (Table 3). These include two different sources that contain information about historical patterns of pedigree linkage or linkage disequilibrium across regions. In all, this is substantially more support for our assembly than was produced by previous Hi-C based assemblies (e.g., [41]; [42]; [43]). Finally, we also note that these independent sources of evidence counter any potential criticism of the fact that our genome assembly (using individual ‘15944’ from the SNPRC baboon colony) comes from a different individual from the previous baboon assembly (individual 1X1155 from the SNPRC baboon colony). In particular, the linkage and linkage disequilibrium based approaches that we used implicitly average across individuals, and make it much more likely that the differences that we observe are not due to polymorphic structural variation in olive baboons.

## METHODS

### Genome Sequencing

#### Index animal

We used individual number 15944 (currently deceased) from the SNPRC pedigreed baboon colony for all of the sequencing and genome assembly work associated with this project.

#### 10x Genomics sequencing

High molecular weight genomic DNA extraction, sample indexing, and generation of partition barcoded libraries were performed according to the 10x Genomics (Pleasanton, CA, USA) Chromium Genome User Guide and as published previously ([28]). An average depth of ~60X was produced and analyzed for this project.

#### Oxford Nanopore sequencing

Libraries for the Oxford Nanopore sequencing were constructed as described previously ([45]) using DNA derived from whole blood. The sequencing was conducted at Genentech, Inc. (South San Francisco, CA, USA); we analyzed data with an average depth of ~15X for this project.

#### Bionano optical maps

High-molecular-weight DNA was extracted, nicked, and labeled using the enzyme Nt.BspQI (New England Biolabs (NEB), Ipswich, MA, USA), and imaged using the Bionano Genomics Irys system (San Diego, CA, USA) to generate single-molecule maps for assessing breaks in synteny between Panu_3.0 and Panubis1.0.

#### Hi-C sequencing

High molecular weight DNA from Jenny Tung (Duke University) was sent to Phase Genomics. ~15X Hi-C data was obtained using previously described techniques [46].

### Linkage disequilibrium analyses

We estimated the scaled recombination rate ρ (= 4Nr where N is the effective population size and r is the recombination rate per generation) using LDhelmet [47] from 24 unrelated olive baboons [48]. We then identified potential breaks in synteny as regions with total ρ > 500 and ρ / bp > 0.2. We considered there to be evidence of a synteny break if one of these regions was within 50 Kb of a potential breakpoint (as identified in Panu_3.0 vs. Panubis1.0 comparisons). The false discovery rate for this definition is ~4%.

The LDhelmet data used a variant call set mapped onto the old assembly Panu_2.0. For the potential breaks in synteny identified above, we used liftover to convert the breakpoints into Panu_3.0 coordinates and verified that Panu_2.0 and Panu_3.0 were syntenic with each other across the breakpoints.

Finally, due to the inherent noise in linkage-disequilibrium based estimates of ρ, the lack of evidence for a synteny break in Panu_3.0 is not positive evidence that the Panu_3.0 assembly is correct.

### Inference of crossovers in a baboon pedigree

We utilized a previously described vcf file for the baboons shown in Figure 6 which was mapped using Panu_2.0 coordinates and lifted over to Panu_3.0 coordinates. We considered only biallelic SNPs, and required a depth ≥ 15, QUAL > 50 and genotype quality (GQ) ≥ 40 in order to make a genotype call. We further required an allelic balance (AB) of > 0.3 for heterozygote calls and AB < 0.07 for homozygote calls, and excluded all repetitive regions as described in [48].

**Figure 6.**
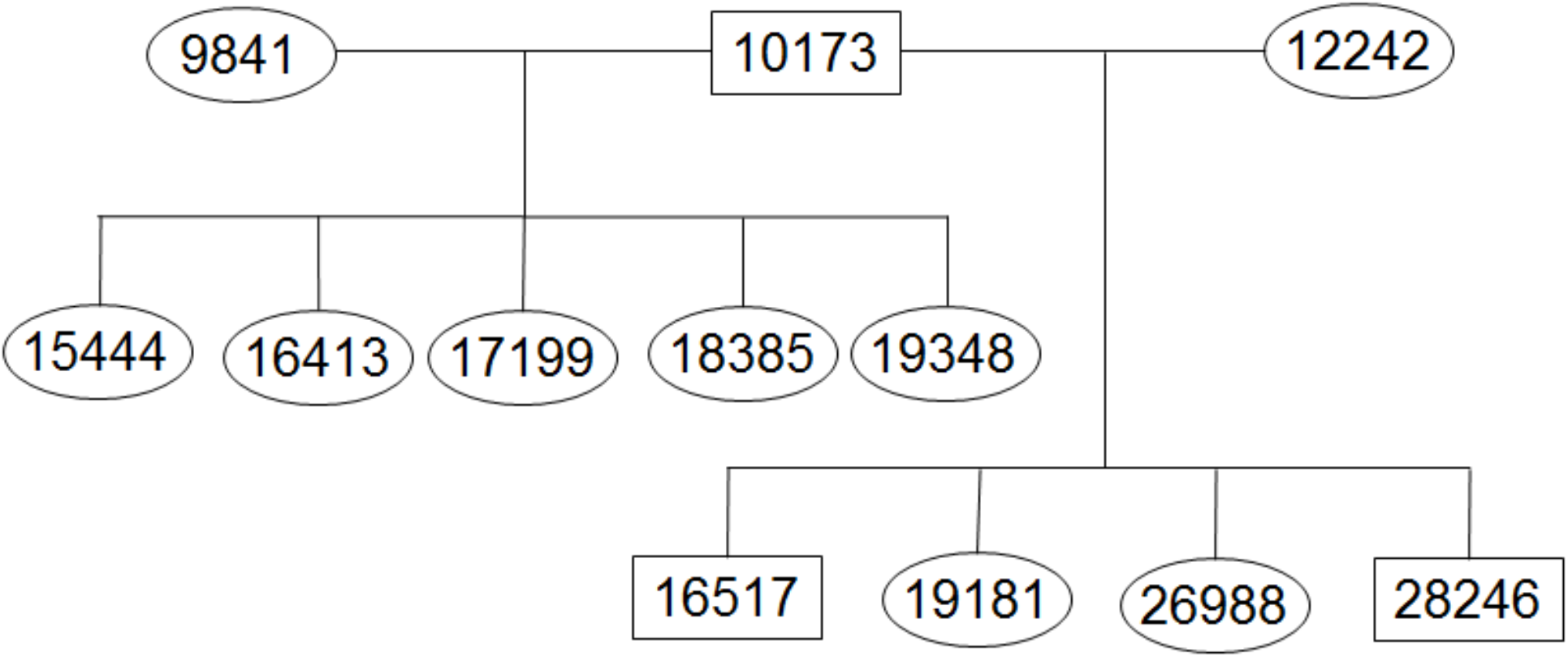
Pedigree of baboons used in linkage analysis.

We focused our analyses on those SNPs that were most informative about recent crossover events. For example, to detect paternal crossovers, we restricted our analyses to SNPs where 10173 was heterozygous, both 9841 and 12242 were homozygous, and all 9 offspring had genotype calls. (For maternal crossovers, we required 10173 to be homozygous and both 9841 and 12242 to be heterozygous.) For these sites, it is straightforward to infer which allele (coded as 0 for reference allele and 1 for alternative allele) was passed on from 10173 to his offspring. While the haplotypic phase of 10173 is unknown, we can infer crossover events based on the minimum number of crossovers needed to be consistent with the observed patterns of inheritance in the offspring of 10173 ([49]). For example, Figure 5c shows that the inheritance pattern near position 29.38 requires at least 3 crossovers (e.g., in individuals 17199, 18385 and 19348).

For each potential error in the Panu_3.0 assembly, we converted the breakpoint location into Panu_2.0 coordinates and verified synteny between Panu_2.0 and Panu_3.0 across the breakpoint region. We then determined whether there were an abnormally large number of crossovers inferred right at the breakpoint. Specifically, if we inferred at least 3 crossover events (out of 18 total meioses, 9 paternal and 9 maternal), then we considered this as evidence that the Panu_3.0 assembly is incorrect, as in Figure 5c (cf. ‘Linkage Support’ column in Table 3). Note that the converse isn’t true: fewer than 3 inferred crossover events is not evidence that the Panu_3.0 assembly is correct at a particular location.

## ACKNOWLEDGEMENTS

We thank Jenny Tung for providing some of the high-molecular weight DNA used in this study. The work was supported in part by NIH grants R24 OD017859 (to LAC and JDW), R01 GM115433 (to JDW), R01 GM094402 (to YSS), R01 HG005946 (to PYK) and by a Packard Fellowship for Science and Engineering (to YSS). YSS is a Chan Zuckerberg Biohub Investigator.

## AUTHOR CONTRIBUTIONS

JDW, LAC and YSS conceived the project. JG, SD, SS, MLS. and PYK generated data for the project. MLS and SSB performed the genome assembly. SSB, MLS, JR and JDW performed the other analyses. SSB and JDW wrote the manuscript with contributions from all authors.

## DATA AVAILABILITY

All of the raw sequence data from individual 15944, as well as the Panubis1.0 assembly are available without restriction from NCBI under BioProject PRJNA527874.

