## Supplementary Information for "Accurate assembly of the olive baboon (*Papio anubis*) genome using long-­read and Hi-C data"

This document contains Supplementary Figures S1-S5.

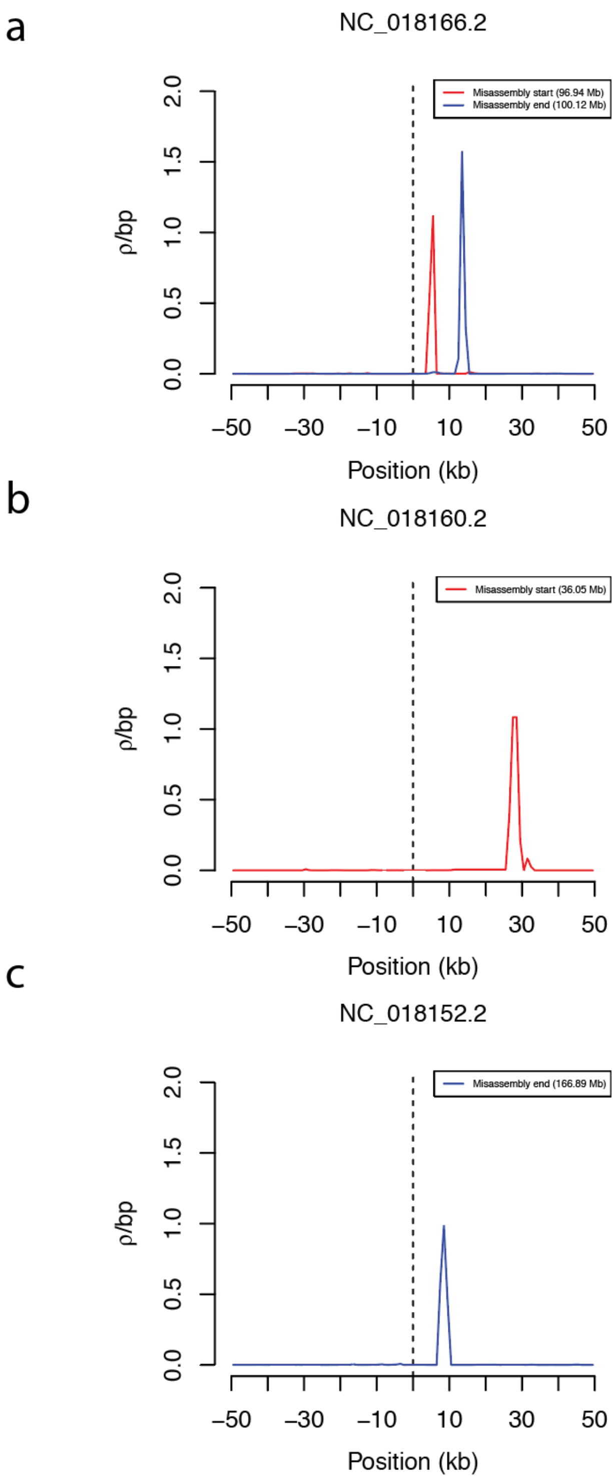


**Figure S1: Linkage disequilibrium based evidence for misassemblies in Panu_3.0.** Estimates of the population recombination rate ρ near the potential synteny breaks of the misassemblies identified in chromosomes a) NC_018166.2 b) NC_018160.2 and c) NC_018152.2. Red represents the beginning of a misassembly event and blue represents the end of a misassembly event.

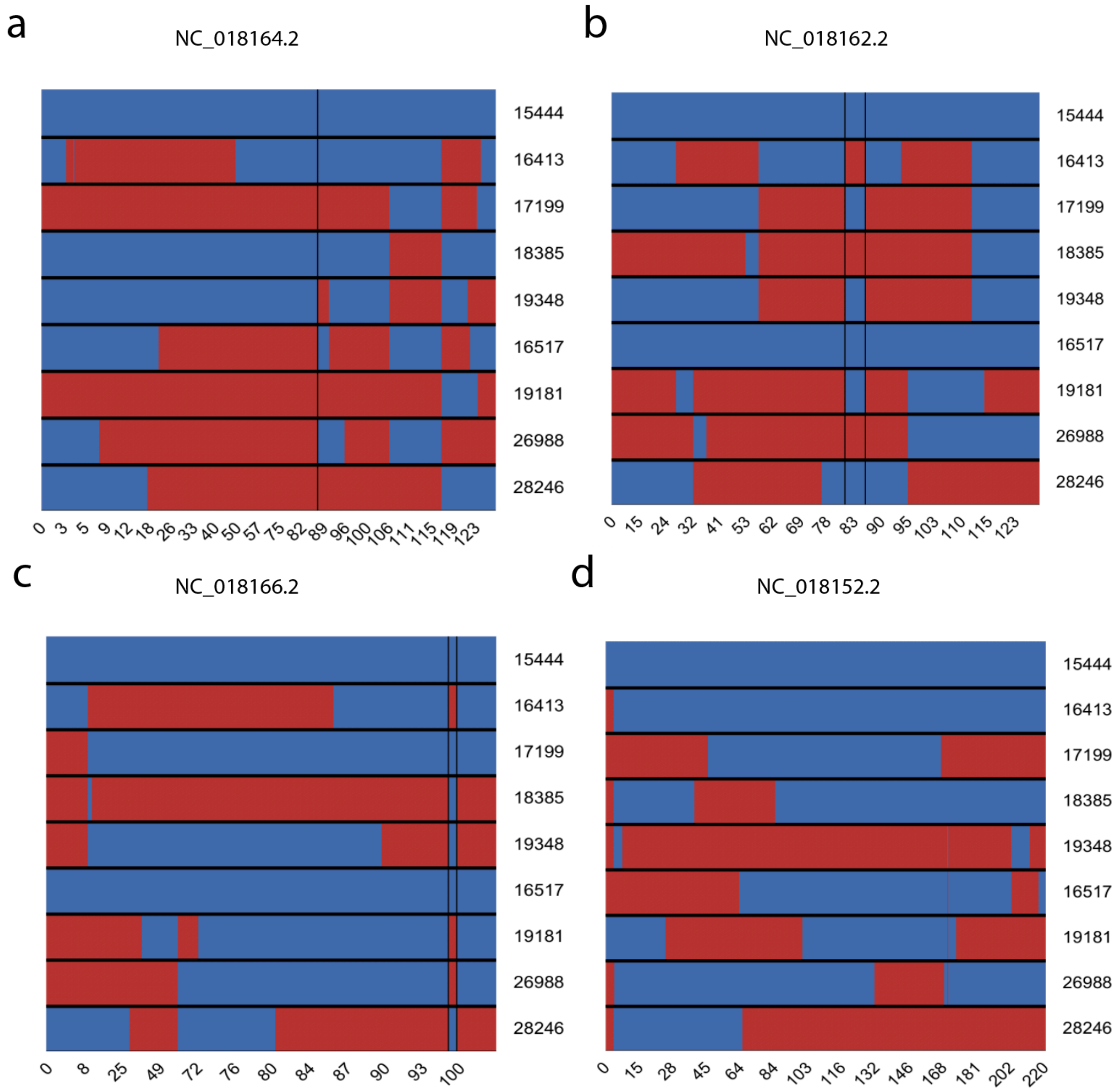


**Figure S2: Recombination based evidence for misassemblies in Panu_3.0.**
Shown on the x-axis is positions along chromosomes in Panu_3.0 where each row represents one of the 9 offsprings of sire 10173. Switches between red and blue within a row represent a recombination event. The vertical black lines represent locations where three or more recombinations occur at the same locus indicating a potential misassembly; except in (d) where recombination occurs at ~167Mb but is not shown by a vertical black line.

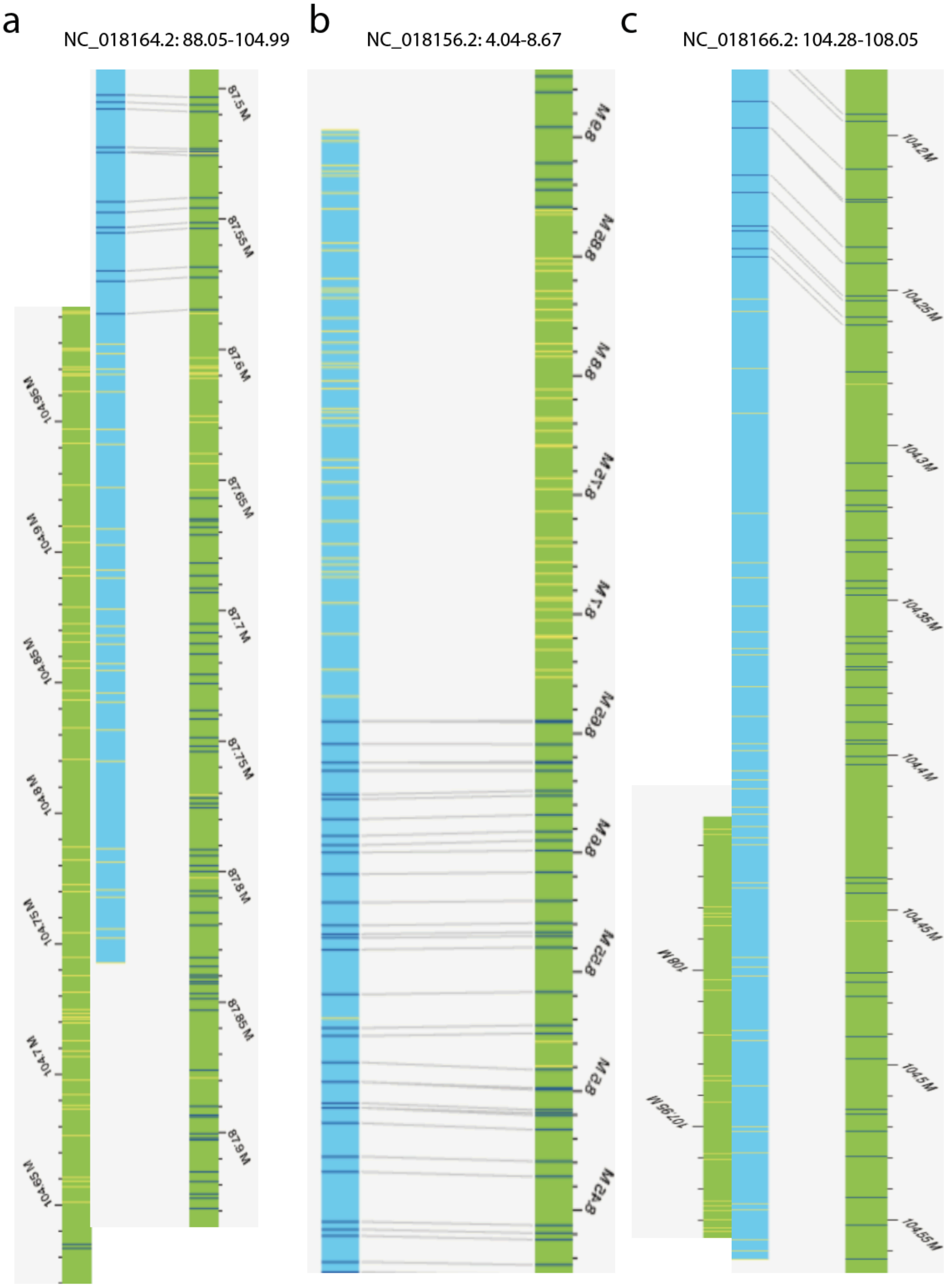


**Figure S3: Evidence for inversions in Panu_3.0 based on bionano alignment.**
a) Inversion on chromosome NC_018164.2 demonstrated by bionano optical map alignment.
b) Alignment to bionano optical map shows inverted coordinates due to an inversion on chromosome NC_018156.2.
c) Bionano optical map alignment shows an inversion on chromosome NC_018166.2.

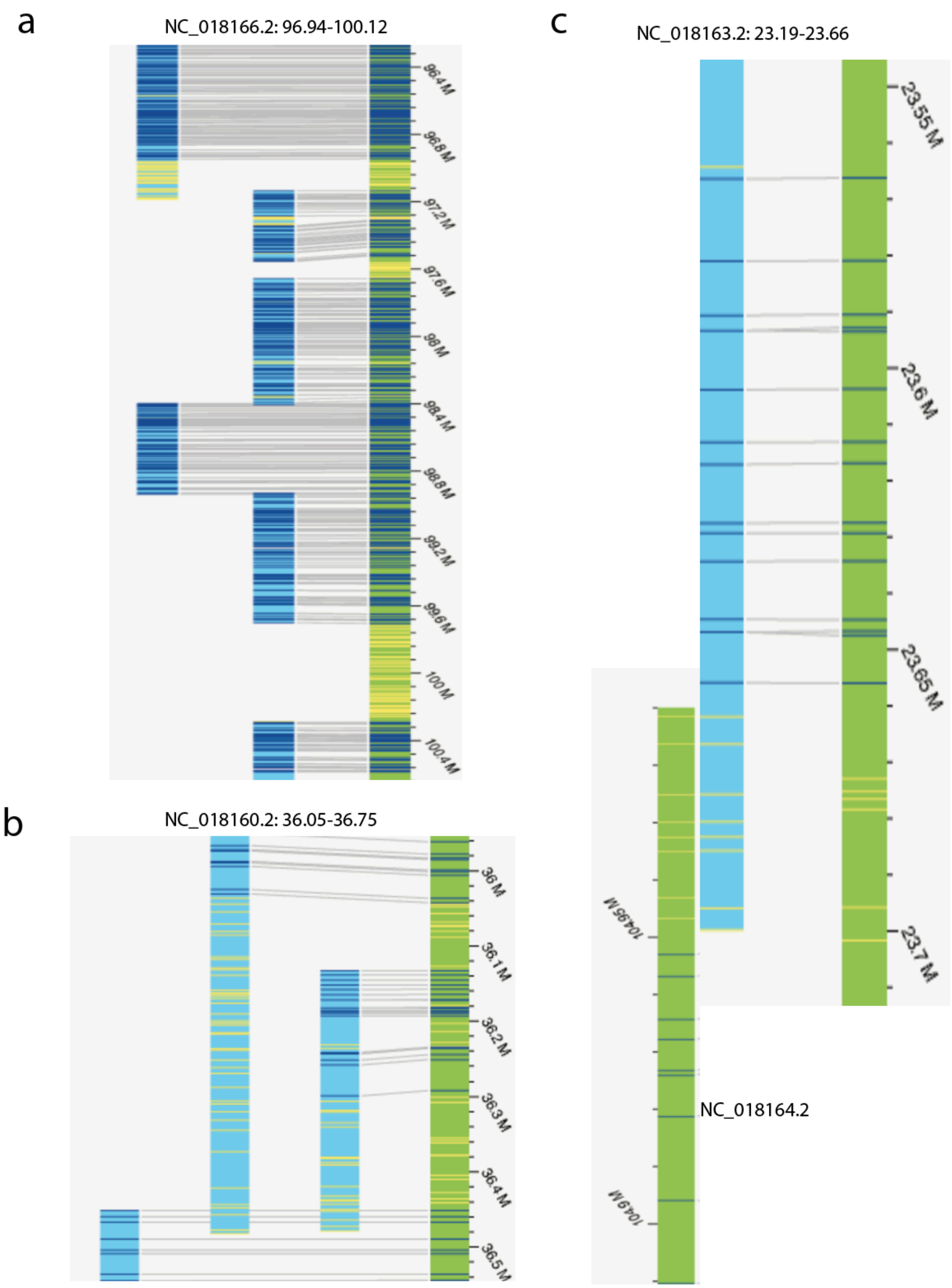


**Figure S4: Evidence for translocations in Panu_3.0 based on bionano alignment.**
a) Breaks in bionano alignment on chromosome NC_018166.2 indicate a misassembly.
b) Bionano optical map alignment demonstrate a misassembly on chromosome NC_018160.2.
c) Bionano optical map alignment shows a translocation between chromosomes NC_018163.2 and NC_018164.2.

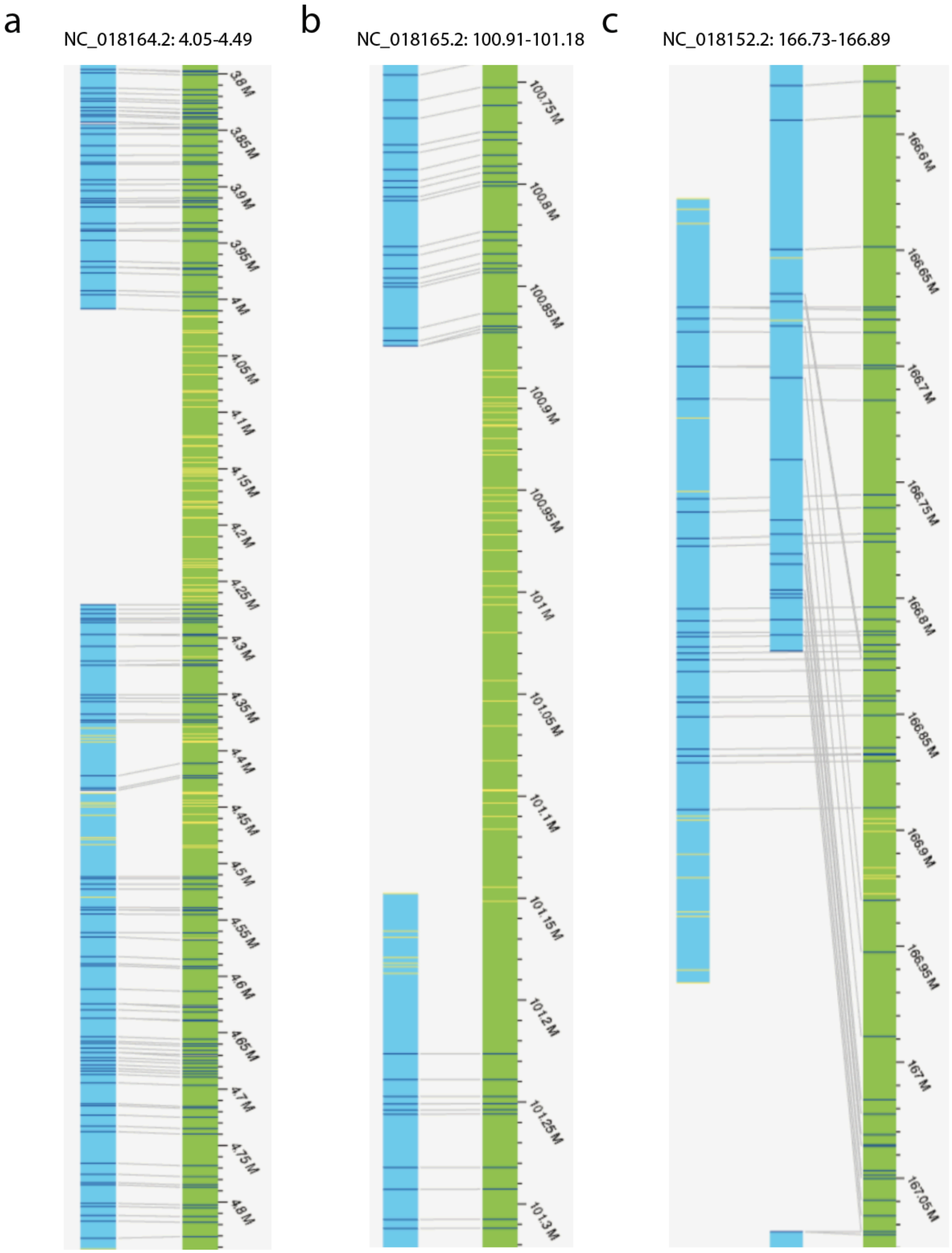


**Figure S5: Evidence for translocations in Panu_3.0 based on bionano alignment.**
a) Breaks in bionano alignment on chromosome NC_018164.2 indicate a misassembly.
b) Bionano optical map alignment demonstrate a misassembly on chromosome NC_018165.2.
c) Bionano optical map alignment shows a translocation on chromosome NC_018152.2.
